# A scalable, multi-wavelength, broad bandwidth frequency-domain near-infrared spectroscopy platform for real-time quantitative tissue optical imaging

**DOI:** 10.1101/2021.07.02.450969

**Authors:** Roy A. Stillwell, Vincent J. Kitsmiller, Alicia Y. Wei, Alyssa Chong, Lyla Senn, Thomas D. O’Sullivan

## Abstract

Frequency-domain near-infrared spectroscopy (FD-NIRS) provides quantitative noninvasive measurements of tissue optical absorption and scattering, and provides a safe and accurate method for characterizing tissue composition and metabolism. However, the poor scalability and high complexity of most FD-NIRS systems assembled to date have contributed to its limited clinical impact. To address these shortcomings, we present a scalable, digital-based FD-NIRS platform capable of measuring optical properties and tissue chromophore concentrations in real-time. The system provides single-channel FD-NIRS amplitude/phase, optical property, and chromophore data at a maximum display rate of 36.6 kHz, 17.9 kHz, and 10.2 kHz, respectively, and can be scaled to multiple channels as well as integrated into a handheld format. The entire system is enabled by several innovations including an ultra-high-speed k-nearest neighbor lookup table method (maximum of 250,000 inversions/s for large 2500×700 table of absorption and reduced scattering coefficients), embedded FPGA and CPU high-speed co-processing, and high-speed data transfer (due to on-board processing). We show that our 6-wavelength, broad modulation bandwidth (1-400 MHz) system can be used to perform 2D high-density spatial mapping of optical properties.

## Introduction

Near-infrared spectroscopy (NIRS) is used to noninvasively characterize the absorption and scattering of light in centimeter thick tissues. This information can be used to assess tissue composition (hemoglobin, water, lipid) (1, 2) and has been applied in clinical and preclinical research for noninvasive brain imaging (3, 4) neonatal care (5), breast tumor imaging (1, 6), muscle training (7), and critical care (8). To date, clinically established NIRS methods are based on continuous-wave (CW) (i.e., time-independent) optical illumination, however time-resolved DOS in both the time-(TD-NIRS) and frequency-domains (FD-NIRS) can provide significant benefits. CW-NIRS assesses changes in light intensity and are therefore largely insensitive to alterations in optical scattering that can occur during the measurement period and between individuals. This limits the quantitative accuracy of CW-NIRS measurements and impedes direct comparison of tissue measurements between individuals and longitudinally over long periods of time.

Alternatively, time-resolved based methods such as FD-NIRS can separate light absorption from scattering, provide more uniform and deeper tissue sampling (9), measure fluorescence lifetime, and are widely recognized as providing substantially greater information content, accuracy, and 3D resolution (2, 9, 10). Their quantitative features allow for absolute comparisons between individuals and longitudinal monitoring of subjects with exceptional accuracy and precision over days to years (11, 12). Scattering parameters can also be isolated and used to provide additional contrast (13). However, despite these advantages, development of clinically-compatible FD- and TD-NIRS instrumentation has been impeded by their poor scalability for high density, high resolution imaging and is relatively complex and slow operation compared to CW-NIRS.

In this work we describe a new FD-NIRS platform that aims to overcome the limitations of scalability and complexity by providing high-speed data acquisition and display of multi-frequency (1-400 MHz) and multi-wavelength (690, 785, 808, 850, 940, and 980 nm) FD-NIRS derived optical properties. Similar to recent work that has demonstrated high-speed FD-NIRS data acquisition (12, 14, 15), we employ an all-digital detection and signal generation scheme with a high-speed analog-to-digital converter (ADC) and a direct digital synthesizer (DDS) to eliminate the slow speed, size and complexity of NA-based systems (16). However, high-speed ADCs have the downside of generating extremely large datasets (in our system around 5.9 GB per second) which is difficult to stream or display in real-time. Because of this, to the best of our knowledge, no FD-NIRS systems have demonstrated real-time spatial mapping of chromophores at display frame rates >60 Hz (1 kHz speed shown in this work), which is critical for providing immediate lag-free feedback to users, especially in a clinical setting. To remedy the shortcoming, we built-in a full digital streaming pipeline and control system inside the hardware which enables real-time chromophore display by using a field programmable gate array (FPGA) and co-processor (CPU), significantly reducing the data streaming output to around 58 bytes. We developed an ultrafast inverse model solution based on a novel (*O*(*log*(*n*)) lookup table approach that is about 1,000x faster than the typical Levenberg–Marquardt algorithm (17) and uses binary trees often deployed in k-nearest neighbor (knn) algorithms (18). In addition, for maximum utility over a wide range of tissue optical properties, our system provides user-selectable single or multi-frequency FD-NIRS modulation.

The platform provides, for the first time, a fully embedded platform for the display of high-speed, high-density 2D quantitative chromophore concentration topography maps–oxyhemoglobin (*HbO*_2_), deoxyhemoglobin (*HHb*), water, and lipid–in real-time. Specifically, the system can display single frequency, single wavelength FD-NIRS amplitude and phase data at a maximum rate of 36,600 Hz, and provides tissue optical property and chromophore concentrations at a maximum of 17,891 and 10,211 Hz, respectively. Metrics for six wavelengths amplitude/phase capture, optical property inversion, and chromophore calculation are 2,560, 2,328 and 2,327 Hz, respectively. Optical property accuracy was compared to a traditional network analyzer-based system (1) and was found to be within 10% over a range of attenuating tissue-simulating phantoms. We show data demonstrating use-case examples of the ultra-high-speed capabilities enabled by our approach including tumor simulating phantom images, as well as in vivo arterial occlusion and pulsatile measurements.

## Methods and Materials

The following section describes the developed system (Section A–D), a brief summary of the optical property recovery methods used (Section E), the characterization measurements undertaken to assess performance and instrument response (Section F), and human in vivo measurement protocols are described in (Section G).

### A. System Overview

Fig.1 shows the overall system design and architecture. The system runs a single or multi-frequency sweep, electrically modulating each of the laser diode sources in a serial manner from an amplified direct digital synthesizer (DDS). Scattered light is collected by a 3mm diameter avalanche photodiode (APD), which is sampled by an analog to digital converter (ADC) at 250 mega-samples-per-second (MSPS). We currently employ sample sizes of 4096, though arbitrary sample sizes (limited by the 128 GB storage in this system) can be used at a cost of lower data collection speed, trading off accuracy with speed. A field programmable gate array (FPGA) captures the ADC data codes and performs a Fourier transform (FT) to extract FD-NIRS phase and amplitude, calibrates the data based on a prior calibration phantom measurement to remove system response, and then calculates optical properties using a novel look-up table approach-all in hardware. Measurements are taken and displayed using a wireless device which communicates with the embedded hardware over peer-to-peer WiFi. The display code was written for Apple’s iOS in Swift, the firmware code was written in C++, while the FPGA elements were written in Verilog.

**Fig. 1.**
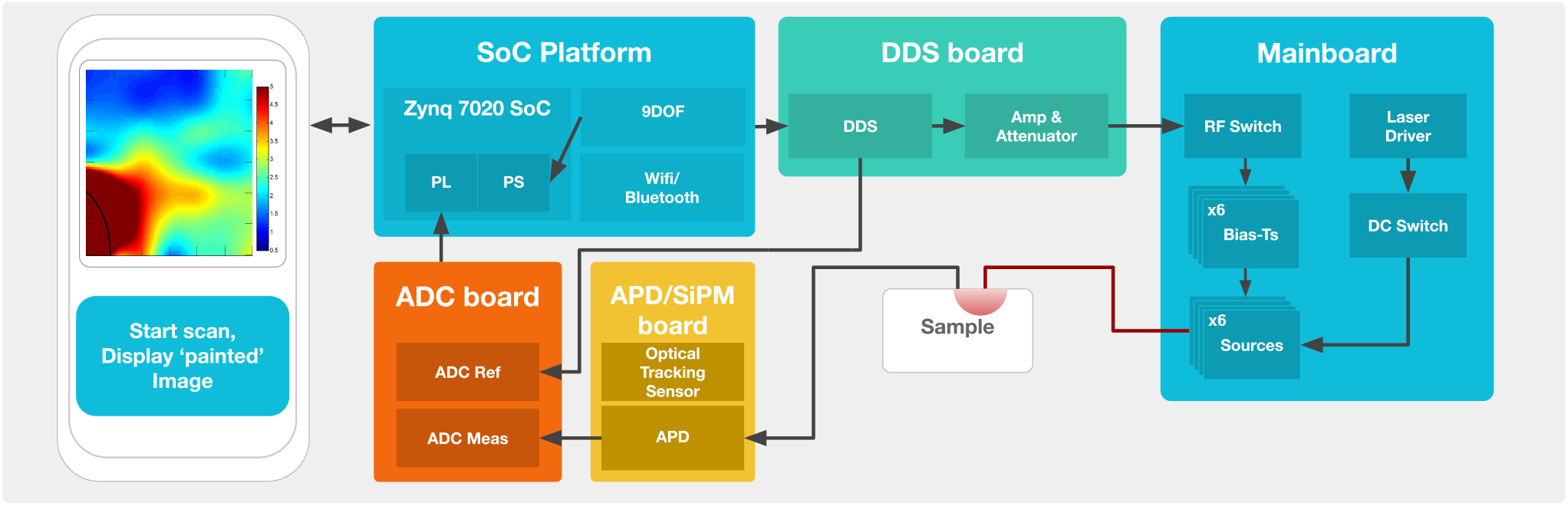
FD-NIRS system design and architecture. Terms: System on a chip (SoC), programming logic (PL), processing system (PS), nine degree of freedom (9-DOF), direct digital synthesizer (DDS), analog to digital converter (ADC), avalanche photodiode (APD).

### B. Signals and Sources

RF from 1-400 Mhz are generated by a DDS chip (Analog Devices AD9912) powered by a high-speed low phase noise 1 Ghz clock (Crystek CCSO-914X-1000), important for precise signal generation. An attenuated portion of RF from the DDS is coupled (Mini-Circuits ZX30-9-4) to the ADC (Analog Devices 9613) as a reference signal, while the remaining is sent to a dynamic gain control chain where a low noise amplifier (LNA; Mini-Circuits TSY-172LNB+) is chained to a digitally controlled attenuator (Skyworks SKY12347-362) with 0 to 31.5dB of attenuation.

The system uses six light sources: five edge-emitting laser diodes (EEL) at 690, 785, 808, 850, and 980 nm (Thorlabs HL6738MG, L785P090, L808P200, L850P030, L980P030), and a vertical cavity surface emitting laser (VCSEL, Vixar) at 940 nm. We designed an aluminum block heatsink to house the sources for temperature and power stability. Each laser is fiber coupled to a 400 *μ*m multimode fiber, with all six fibers being collected into a single ferrule. A single laser driver (Thorlabs MLD203CHB) is connected to two identical 1×3 DC switches (Analog Devices ADG802BRTZ) on the mainboard (Fig.1) providing 6 DC outputs. Each output provides a DC bias and is connected to each source with a bias tee, while the RF for each source is switched with an SP8T RF switch (Analog Devices HMC253AQS24E).

### C. Detection

Light is collected with a 3mm avalanche photodiode (Hamamatsu S2384) that directly touches the sample surface. The detected signal is amplified +15 dB with an LNA (Mini-Circuits TSY-172LNB+) to allow for full use of the full-scale voltage (FSV) of the ADC, and can be bypassed for close source/detector separations so as not to exceed FSV of the ADC. Typical signal levels from the APD range from 0 to −70 dBm, and thus fit within the ADC’s FSV range to maximize ADC resolution. The single ended signal from the APD is then converted into a differential signal and ac coupled to the ADC. The ADC samples at 250 MSPS at 12-bits of resolution, dual channel, and with 1.75V FSV. The ADC outputs digital codes to the FPGA in the system on a chip (SoC). Modulation frequencies above 125 Mhz are aliased (due to the 250 MSPS sampling rate), however since all modulation frequencies are known, aliased frequencies in overlapping Nyquist zones are removed (16).

### D. Signal Processing and Control

12-bit data codes from the ADC encode the voltage levels for two channels: the measured APD channel and a reference DDS channel that serves as the amplitude and phase reference (shown in Fig. 1). The ADC 12-bit codes are processed in hardware using an FPGA/Co-processing chip (Xilinx Zynq 7020) facilitated by a commercial off the shelf (COTS) SoC platform (Krtkl Snicker-doodle Black). The system runs Linux in the processor system (PS), which then controls the data processing being done in the programming logic (PL). We developed a custom high speed TCP/IP data packet protocol that connects the mobile display device wirelessly to the system using AES encrypted peer-to-peer Wi-Fi.

The signal processing chain is shown in Fig. 2. In the pre-processing steps (done immediately after a user sets the measurement parameters in the GUI), the coefficients for the Goertzel algorithm (described in detail in the following paragraphs) are calculated on the device. Dual-channel raw ADC sample codes are captured by the FPGA and as each sample is received, they are pipelined to the Goertzel Algorithm block which collects and converts the samples to real and imaginary values for each frequency used in a measurement as shown in Fig. 2. Of note is that with this pipeline method, the finished Fourier transform output of amplitude and phase values are ready within a single clock of the ADC sampling being finished, allowing for immediate use of the FT data. The measured signal is then calibrated with data from a calibration phantom to remove the internal system response.

**Fig. 2.**
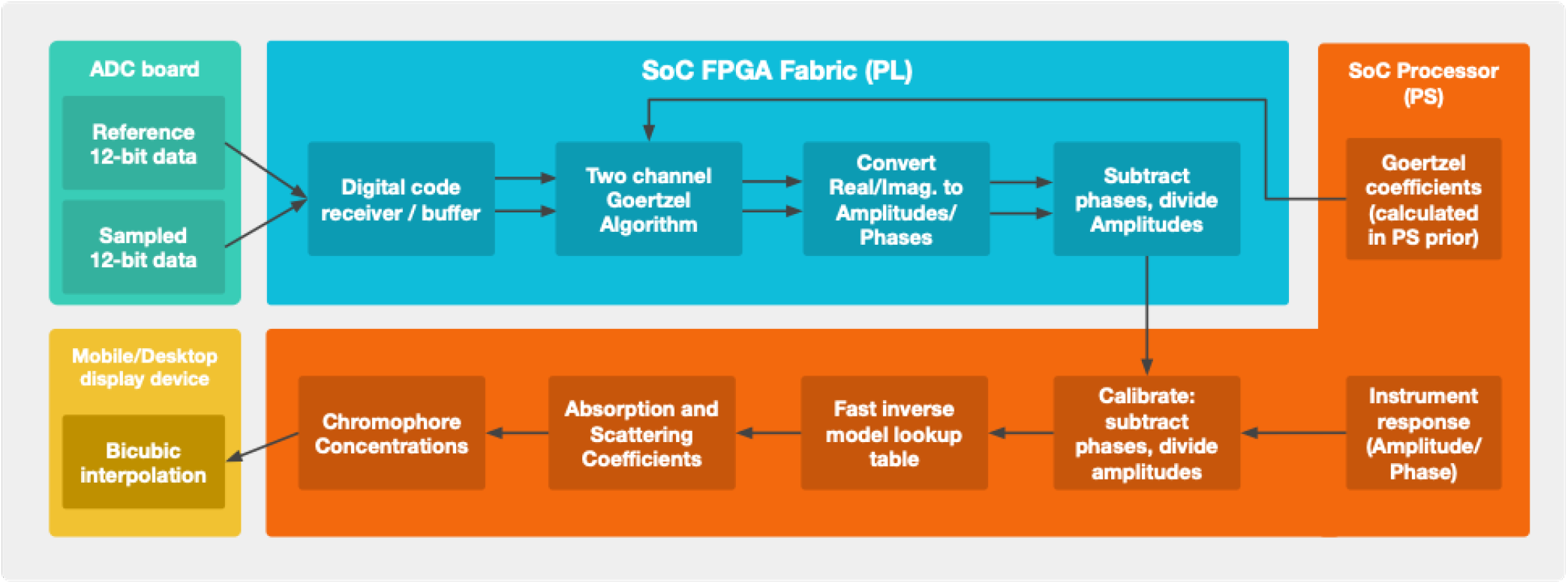
Signal processing block diagram.

Raw digital data codes from the ADC are processed directly within the FPGA fabric. A typical approach(16) might use a fast Fourier transform (FFT) algorithm. However, because we need data from only one of the frequency bins at a time, we can significantly reduce computation time by using an optimized discrete Fourier transform (DFT). An algorithm invented by Goertzel(19) computes the *k*th DFT portion of a signal *x*[*n*] of length *N*. Essentially, the Goertzel algorithm performs a computation of a single DFT coefficient. This can be advantageous over a full DFT for several reasons. When only a few values of the spectral components of a signal are required, rather than the whole spectrum, the algorithm can be significantly faster. Because one can consider the Goertzel as an infinite impulse response (IIR) filter(19), the computation can begin immediately at the arrival of a sample, a key to real-time processing. This is opposed to an FFT where the entire sample set is required first before computation. Additionally, there is no need to output data in bit-reverse order or build a re-order logic block in FPGA fabric. As such, an extremely low latency (single clock cycle) pipeline can be used to do the computation. Furthermore, FFT efficiency is dependent largely on the sample length, *N*, which is most effective as a power of two. For the Goertzel algorithm, any arbitrary N size is possible without adding computational complexity. While others have used the Goertzel algorithm for processing FD-NIRS signals (15, 20), our approach differs in that the constants used in the Goertzel algorithm are pre-calculated and stored in registers in the FPGA for each frequency in a sweep, where others have only done single frequency calculations in an FPGA, or a multi-frequency Goertzel algorithm done less efficiently in software. The allows for single-clock computation for each frequency used.

The Goertzel approach minimizes the number of calculations required for a real-valued input signal. For an input signal, 2*N* additions and *N* multiplications are performed, resulting on the order of 3*N* +2 computations for each frequency. This approach is more advantageous as compared to an FFT when the number of frequency bins *K* < 2*log*2*N*. Memory requirements are also reduced when *K* < 4/7*N* as compared to an FFT (19). In our case *K* is simply one (a single frequency bin). As an example, for a sample length of 65,536, the above approach achieves a 32x decrease in computation and a 37,000x decrease in required memory usage over an FFT.

During an FD-NIRS measurement the system will generate large quantities of data for a single measurement (16 kB at 4096 samples, 1*λ,* 1*f*) or around 8 gigabits-per-second (sampled at 250 MSPS). Streaming this much data in real time to an external device is not practical. Our approach reduces this data output by embedding the FT and recovery of optical properties into the hardware, reducing the data output to only 8 bytes per measurement (1*λ,* 1*f*) which is easily transferred at a real-world 2 megabits-per-second wireless bandwidth.

### E. Model Inversion and Chromophore Estimation

Raw amplitude and phase data captured in the signal processing chain as described in Section D represent a convolution of attenuation and phase delay from the measured tissue and the instrument response. To remove instrument response, data is calibrated against a silicone phantom with known optical properties (10) and the data is stored in on-board memory. Pre-calculated inverse solutions to the p1 semi-infinite model (21, 22) are then loaded into memory as lookup-tables for high-speed optical property recovery. There is a table for each frequency in the sweep and for each source detector separation (SDS) used. The values are stored as complex and each table spans *μ_a_* and *μ_s_* values. The tables are indexed into binary search trees to enable very fast searching on the order of *Olog*(*n*) (18). After the Goertzel algorithm calculates the measured real/imaginary data for each channel (in parallel to the device sampling from the ADC), the system completes a k-nearest neighbor (knn) search (18) for a *μ_a_*, *μ_s_* solution that matches the measured complex value, where k is the number of dimensions (in this case two, for the real and imaginary part). For multi-frequency sweeps, these *μ_a_*, *μ_s_* solutions are averaged to find final *μ_a_*, *μ_s_* values. The results of the knn lookup table method are covered in detail in the following sections. Chromophore concentrations are then calculated in firmware using the known extinction coefficients (23–25).

### F. Characterization Methods

The measurements we performed to characterize the FD-NIRS platform performance are presented in this section. Unless specified otherwise, all wavelengths of 690, 785, 808, 850, 940, and 980 nm were used. The sources were modulated at a single frequency of 70 MHz (though any frequency from 0.1-400 MHz could be used). All measurements were collected in a reflection geometry at an SDS of 23 mm. 4096 samples were taken at a speed of 250 MHz with the ADC. Optical and chromophore properties were extracted using the method described in Section E. The table size used for recovery was 2500 × 700 (*μ_a_,μ_s_*) as we found this table size brought the quantization error to below 1% for both *mu_a_* and *mu_s_*. Optical power was set to 20 mW for all sources as measured from the fiber ferrule.

#### F.1. Noise Characterization

The noise floor of the system was measured in two ways. First, a ‘dark’ measurement was taken wherein the 3mm APD detector in the system was biased and optically blocked, while the laser source power systems were energized. The electrical noise floor was also measured without the APD or laser source power supplies energized, which should result in a much lower noise floor as optical systems are typically detector limited. Samples were taken with the ADC at 4,096 and 65,536 samples for the dark measurement, and at 65,536 for the electrical measurements to accurately measure the extremely low noise floor of the system (26).

#### F.2. Accuracy and Precision

A series of 10 solid silicon tissue simulating phantoms were prepared with varying *μ_a_* from 0.002-0.020 cm^−1^ and *μ_s_* varied from 0.5-1.4 cm^−1^ (27). Optical properties for the system were recovered using the p1 semi-infinite model in the form of a lookup table as described in Section E and calibrated with a silicone phantom with known optical properties. The gold standard system was VNA based as described in (1) and was used to measure the phantoms with a self-calibrated multi-distance method (SDS ranging from 1.5 to 3.0 cm) (22, 28). To assess precision, 12 seconds of data (30,720 samples for all six sources; 2,560 Hz display rate, 70 MHz single frequency) was collected on a tissue simulating phantom (designed for *μ_a_*= 0.01 mm^−1^ *μ_s_*= 1.0 mm^−1^ at 660 nm).

#### F.3. Inversion Speed

Inversions per-second measurements were made on two different hardware schemes. The execution speeds were averaged over 10,000 look-ups, single threaded, and measured using on-board system clocks, accurate to around 1 *μ*s. The ‘Modern CPU’ system used in Section J was a typical workstation processor (AMD 3960X, 3.8 GHz), where as the ‘SoC’ was the system on a chip (Zynq 7020, Xilinx).

#### F.4. 1D Line Scans and 2D Imaging with Tissue-Simulating Phantoms

To demonstrate high-density 1D line scans and 2D imaging, a series of solid silicone tissue simulating phantoms were prepared with a simulated tumor inclusion 1 mm below the surface with diameters of 1.0, 1.5, 2.0, and 3.0 cm (substrate: *μ_a_*= 0.005-0.008 *mm*^−1^, *μ_s_*= 0.6-0.9 *mm*^−1^; inclusion *μ_a_*= 0.013 *mm*^−1^, *μ_s_*= 1.4 *mm*^−1^ at 660 nm). The probe housing the optical ferrule and 3 mm APD was moved across the surface over the phantom and tumor inclusions at a constant rate. 2D images of the 30 mm diameter inclusion phantoms were created using high-density vertical and horizontal data. All 6 sources were used. The line scans were combined into a single matrix, after which bi-cubic interpolation was employed to generate the image.

### G. In Vivo Measurements

In vivo arterial occlusion measurements were obtained by placing a blood pressure cuff on the upper arm of a healthy 39 year old male volunteer. The human study was approved by the Institutional Review Board of the University of Notre Dame and the participant provided written informed consent. Continuous measurements acquired with the system were displayed and recorded in real-time at 2.6 kHz with all 6 sources. Optical properties were recovered following the method outlined in E. The volunteer began the experiment in a quiet room at rest for 20 minutes prior to measurement. Data was collected on the right forearm for 2 minutes at rest, with the volunteer breathing normal and relaxed (baseline). After 2 minutes, the cuff was inflated to 250 mmHg to restrict blood flow (occlusion). After 5 minutes of occlusion, the cuff was slowly released, after which data was recorded for another 5 minutes (recovery).

An extremely high-speed pulse measurement (5.4 kHz) was also taken on the right-wrist of the same volunteer. Data was acquired for 35 seconds with two wavelengths (690 nm and 850 nm) to recover *HbO*_2_ and *HHb* concentrations. Data was smoothed using a 100-pt moving average. For this measurement the SDS was set to 12 mm as the radial artery targeted was near to the surface of the skin. Prior to the measurement the volunteer was at rest for 20 minutes in a quite room. During the measurement the volunteer was breathing normally. The volunteer’s heart rate was simultaneously measured at around 67 BPM via a manual palpation].

## Results

### H. System Characterization

The six light sources are capable of maximum optical powers ranging from 30-200 mW. The DC and RF switching integrated circuits can operate very fast (~90 ns), but a typical laser settling time was found to be 1 *μ*s for this system, limiting source switching speed to around 1 MHz. The DC laser driver exhibits excellent stability with 12 *μ*A average root-mean-square current with *μ*W power stability. The laser driver can deliver up to 200 mA with 3V compliance voltage and the DDS can deliver 0-13.7 mA, with a maximum 7dBm power into a 50 Ω load at 50 Mhz. After the amplifier chain, the maximum RF output was just below 20 dBm.

The overall noise floor of the system was found to be −68 dBm at 4096 and 65,536 samples, suggesting it is the true noise floor. The noise floor data is shown compared to a VNA based gold standard system in Fig. 3 (1). The electrical noise floor, measured without the detector was −93 dBm, illustrating that system sensitivity is limited while the source and detector circuitry is energized. The modulation efficiency during measurements was found to be between 90-95%, depending on the source.

**Fig. 3.**
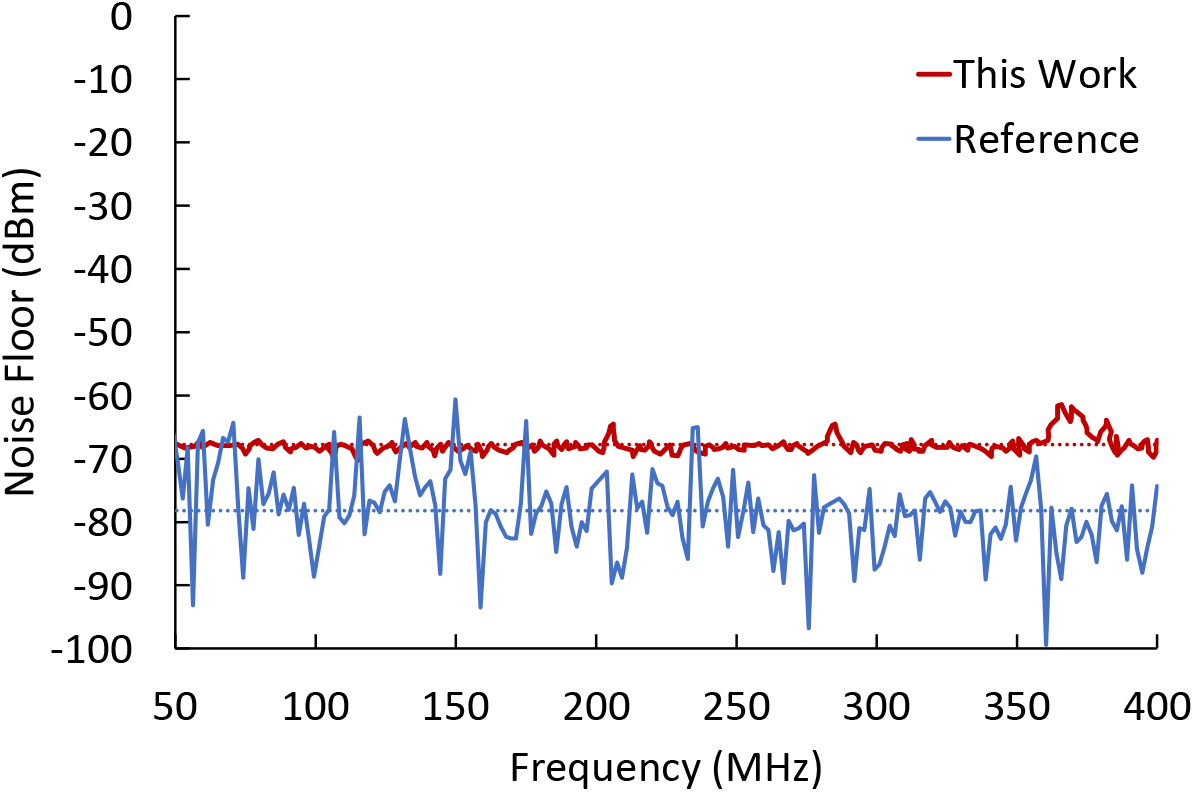
Noise floor of the FD-NIRS system compared to a reference VNA-based system.

### I. Accuracy

Figure 4 shows optical property recovery of a range of tissue simulating phantoms, comparing our system with a gold standard VNA system. We find that for nearly all absorption and scattering values, the system was within 10% of the gold standard across all wavelengths, suggesting acceptable agreement between the systems. *μ*^′^_s_ values had the most agreement with the reference system when values were near 0.75 mm^−1^, while *μ_a_* were in closer agreement with values lower than 0.013 mm^−1^. The mean difference for both *μ_a_* and *μ*^′^_s_ were nearly identical to the VNA based gold standard.

**Fig. 4.**
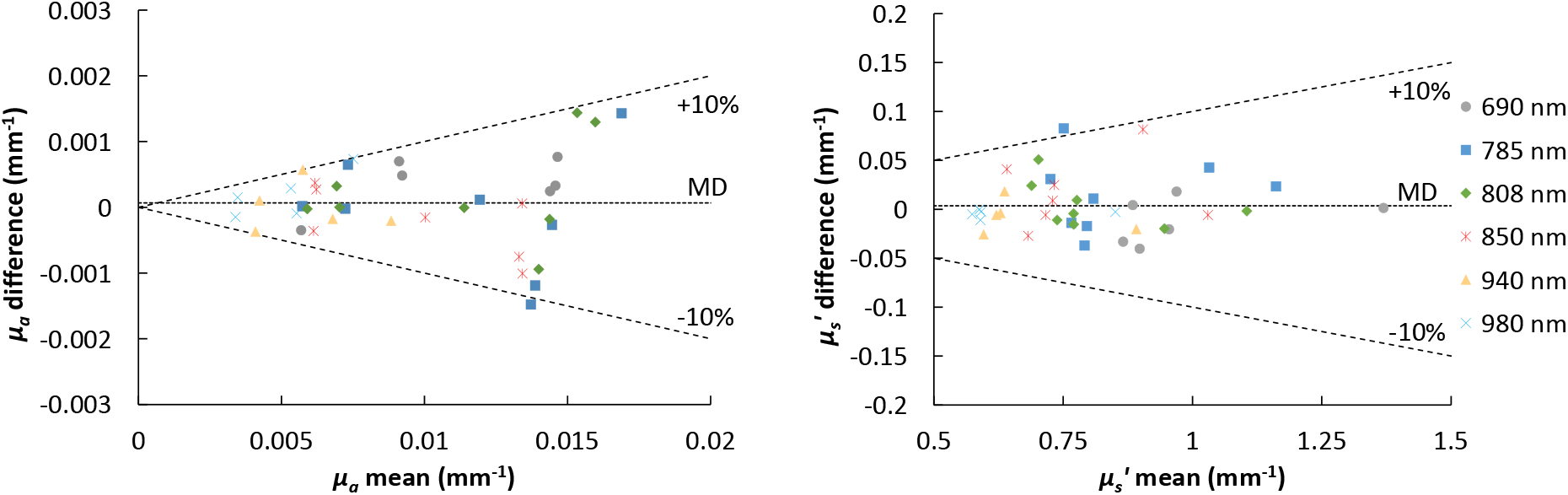
Bland-Altman plots of optical properties measured with the system as compared to the reference VNA based system. The center line shows the mean difference, while the top and bottom dashed lines denote 10% deviation from the mean difference. Nearly all absorption and scattering differences fall within 10% of the VNA gold standard system.

### J. High-Speed Inversion

Table 1 shows the inversion rates of single and multi-frequency optical property recovery using the high-speed knn lookup table method outlined in Section E. A low and high density lookup table for *μ_a_* and *μ_s_* was chosen to show the (*O*)*log*(*n*) (18) scaling made possible with the knn method as values of n (table size) grows. As an example, a 16x increase in table size from 500×500 to 2500×700 only had a 2% decrease in inversion/s speed. The knn lookup table algorithm running on the ‘modern cpu’ (AMD 3960X, 3.8 GHz) is between 17-18x faster in inversion/s over the lower performance SoC (Zynq 7020, 866 MHz) in both single and multi-frequency inversion schemes, as was expected. The knn lookup table method for inversions showed between 170 to 11,000x improvement in speed using modern desktop processing hardware (AMD 3960X, 3.8 GHz) over any system we know of. We also measured between 12 and 1,900x improvement in inversions/s when using the SoC.

**Table 1.** High-speed knn lookup table inversions/s

| Table Size, Frequencies | Modern CPU (Hz) | SoC (Hz) |
| --- | --- | --- |
| 500x500, 1f | 255,916 | 18,772 |
| 2500x700, 1f | 250,771 | 17,841 |
| 500x500, 51f | 46,919 | 2,691 |
| 2500x700, 51f | 11,193 | 1,911 |

### K. High-Speed Acquisition

The device acquires high speed FD-NIRS measurements in a serial manner; each source is modulated sequentially and can be swept in modulation frequency from lowest to highest for multi-frequency measurements. The raw data capture speed of the 250 MSPS ADC dictates the maximum obtainable speed as we need to capture enough of the waveform to extract accurate amplitude and phase information. For example, at a 4096 sample rate, data acquisition takes around 16.4 *μ*s, leading to an upper speed limit of 61,035 Hz (the fastest speed at that sample size). Smaller sample lengths can be used, though at the expense of phase resolution and thus accurate optical property recovery (16). In practice, we found that 4096 is a good balance between data acquisition speed and accuracy of recovered optical properties.

Since our primary aim is to use the instrument to display data in real-time for use in a clinical system, we report our realized display rates for each stage of the data processing and display in Table 2. For a single-source/single-frequency measurement, the maximum rate for amplitude and phase measurement and display is 36,608 Hz, optical properties were found to be measured and displayed at 17,891 Hz, and chromophores at 10,211 Hz. Amplitude and phase from all six sources can be displayed at 2,560 Hz, optical properties at 1,615 Hz, and chromophores at 2,327 Hz. However, this can be further improved as there is a small delay introduced by the serial peripheral interface (SPI) communication driver associated with the switching hardware. A six-source, multi-frequency sweep from 50-300 Mhz at 7 Mhz steps (35 frequencies) results in amplitude/phase display at 175 Hz, with optical properties and chromophores displayed at 60.2 Hz. If ultra high speed data display is not required, such as for single SDS configurations or applications requiring long-term continuous measurements, SNR can be improved by averaging the data. Table 3 illustrates how averaging improves the phase precision at the expense of speed. The standard deviation decreases by more than 8.5x from 0.204 degrees (no averaging) to 0.024 degrees (30,000 samples averaged).

**Table 2.** Display rates in Hz for each phase of the pipeline required to display four chromophore values (*HHb*, *HbO*_2_, *H*_2_*O*, and lipid).

| Parameters | Raw samples | Amp/Phase | OPs | Chromophores |
| --- | --- | --- | --- | --- |
| 1λ, 1f | 61,035 | 36,608 | 17,891 | 10,211 |
| 6λ, 1f | 10,173 | 2,560 | 2,328 | 2,327 |

**Table 3.** Block averaging showing effect on phase precision. 70 MHz, 785 nm source (others wavelengths were similar).

| Sample Blocks Averaged | Acquisition Time(s) | Effective Rate (Hz) | Phase std (°) |
| --- | --- | --- | --- |
| 1 | 0.000027 | 36600.00 | 0.204 |
| 5 | 0.000137 | 7320.00 | 0.108 |
| 10 | 0.000273 | 3660.00 | 0.088 |
| 30 | 0.000820 | 1220.00 | 0.070 |
| 300 | 0.008197 | 122.00 | 0.030 |
| 3000 | 0.08 | 12.20 | 0.026 |
| 30000 | 0.82 | 1.22 | 0.024 |
| 300000 | 8.20 | 0.12 | 0.010 |

### L. Stability and Precision

System precision and stability was assessed by analyzing data collected continuously over a short (12 s) and long (1 hr) periods. For the short-term precision assessment (Table 4), 12-seconds of high-speed data was collected on a tissue simulating phantom. The system displayed a coefficient of variance (CV) of less than 1% for all amplitudes and phase standard deviation typically between 0.21-0.33 degrees, though the 808 nm was slightly higher at 0.56 degrees due to its lower signal strength. A one-hour stability measurement was taken, wherein the same tissue simulating phantom was measured continuously at 2 second intervals. Optical properties were extracted using the method described in Section E. The sources showed good optical property stability for *μ_a_* and *μ_s_* with an average CV of 3.66% and 1.92%, respectively, across all wavelengths as shown in Table 5. We note that the data sampled at 2 Hz was not averaged. If increased precision is necessary, the platform is capable of averaging optical properties for all six wavelengths over longer time periods, as was shown in Table 3.

**Table 4.** High-speed system stability during a 12-second measurement with a display rate of 2,560 Hz, for all six sources at 23 mm and 70 MHz on a tissue simulating phantom (*μ_a_*= 0.01 *mm*^−1^1 *μ_s_*= 1 *mm*^−1^ at 660 nm).

| $\lambda$ (nm) | Amp. std (dbFS) | Amp. CV (%) | Phase std (°) |
| --- | --- | --- | --- |
| 690 | 0.040 | -0.25 | 0.227 |
| 785 | 0.030 | -0.30 | 0.209 |
| 808 | 0.087 | -0.42 | 0.557 |
| 850 | 0.048 | -0.24 | 0.321 |
| 940 | 0.052 | -0.39 | 0.332 |
| 980 | 0.080 | -0.74 | 0.320 |

**Table 5.** Standard deviation and coefficients of variation for optical properties over a 1-hr drift test across the six wavelengths used.

| $\lambda$ (nm) | $\mu_a$ mean / std ($\text{mm}^{-1}$) | $\mu_a$ CV (%) | $\mu_s$ mean / std ($\text{mm}^{-1}$) | $\mu_s$ CV (%) |
| --- | --- | --- | --- | --- |
| 690 | 0.0090 / 0.00014 | 1.57 | 0.96 / 0.0094 | 0.98 |
| 785 | 0.0072 / 0.00018 | 2.50 | 0.80 / 0.0107 | 1.34 |
| 808 | 0.0070 / 0.00037 | 5.29 | 0.77 / 0.0231 | 2.99 |
| 850 | 0.0063 / 0.00041 | 6.48 | 0.72 / 0.025 | 3.50 |
| 940 | 0.0043 / 0.00012 | 2.91 | 0.63 / 0.0085 | 1.35 |
| 980 | 0.0035 / 0.00011 | 3.21 | 0.59 / 0.0079 | 1.34 |

### M. High-density, high-speed measurements of phantom inclusions

#### M.1. 1D Line Scans

Fig. 5 shows representative high density 1D line-scan optical property measurements of tissue simulating phantoms containing optical inclusions. The 808 nm data is shown, though the other five wavelengths had exhibited a similar shape. Data was acquired at full speed of 2.6 kHz, then 30 pt moving averaged, totaling 82,000 views and 492,000 total samples (for all six sources) for each line scan. Capture time for each measurement was just over 12 seconds. The 1/*e*^2^ values (a common beam diameter definition (29)) were calculated for each resulting curve for *μ_a_* and (*μ_s_*) at 1.0, 1.5, 2.0, and 3.0 cm diameters, resulting in errors of 81% (0.4%), 20.6% (0.6%), 8.5% (−1%) and 1.6% (3.3%), respectively. Due to the extremely high density of data points, the curves show how smooth and information rich the data is when using high-speed data acquisition (video available with online version). The data suggests 1/*e*^2^ values of the high-spatial resolution scattering information allows for an accurate estimate on the size of the underlying inclusion at these depths and SDS, but localization in the diffuse optical regime is a more complicated function of SDS, inclusion depth, and optical properties. We also found that high resolution scattering 1/*e*^2^ values better estimated the size of the underling inclusion when compared to low resolution (single cm) sampling. However, the choice of width metric is somewhat subjective (30). Regardless, the data suggests that scattering information may give more accurate localization than absorption as others have suggested (31, 32), but more inquiry is required.

**Fig. 5.**
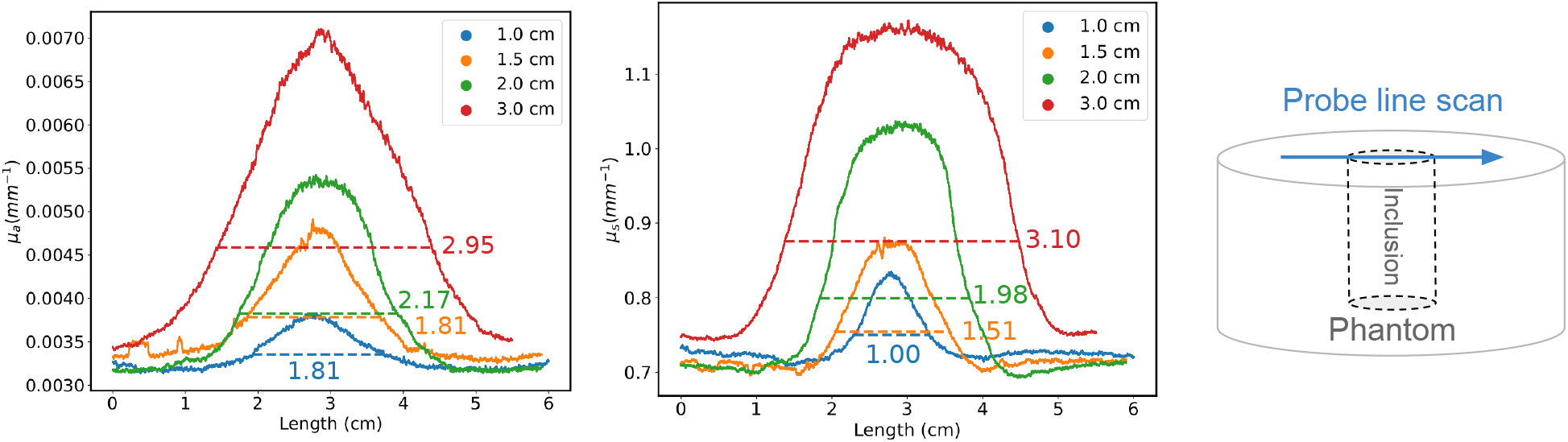
1D line-scans showing (left) absorption and (middle) scattering coefficient data taken of tissue simulating phantoms (substrate: *μ_a_*= 0.005-0.008 *mm*^−1^, *μ_s_*= 0.6-0.9 *mm*^−1^; inclusion *μ_a_*= 0.013 *mm*^−1^, *μ_s_*= 1.4 *mm*^−1^ at 660 nm), shown at 808 nm, single frequency (70 MHz). Simulated tumor inclusions were 1 mm below the surface in the center of the phantom on 10, 15, 20, and 30 mm diameter sizes. (right) Experimental setup diagram.

#### M.2. 2D Spatial Mapping

2D images of the 30 mm diameter inclusion phantoms were created using high-density vertical and horizontal data (8 line scans in each direction) as shown Fig.7c. The high-density data set was captured at a 2.6 kHz data rate, at 22,000 samples per line scan for a total of 352,000 points per image, 2,112,000 total samples for all six sources. The image acquisition time was about 3 minutes. High-density images for all six wavelengths are shown in supplemental Fig. S1. For comparison Fig. 7a shows an image downsampled to 1 cm spacing (comprising of 49 points), which was the measurement spacing used in the ACRIN 6691 DOS imaging trial of breast cancer neoadjuvant chemotherapy response (12). Data from Fig.7a was then interpolated to form Fig.7b. The lower spatial density acquisition image of Fig.7a and Fig.7b suggests valuable absorption and scattering information is missed, as compared to the high-density images in Fig.7c where the 30 mm tumor topography is more accurately represented. This represents highest density 2D topographical FD-NIRS image created to date, with 43,100x times more spatial density than that used in ACRIN 6691 (12) and as reported more recently(33), about a 1,550x improvement in chromophore mapping speed.

**Fig. 6.**
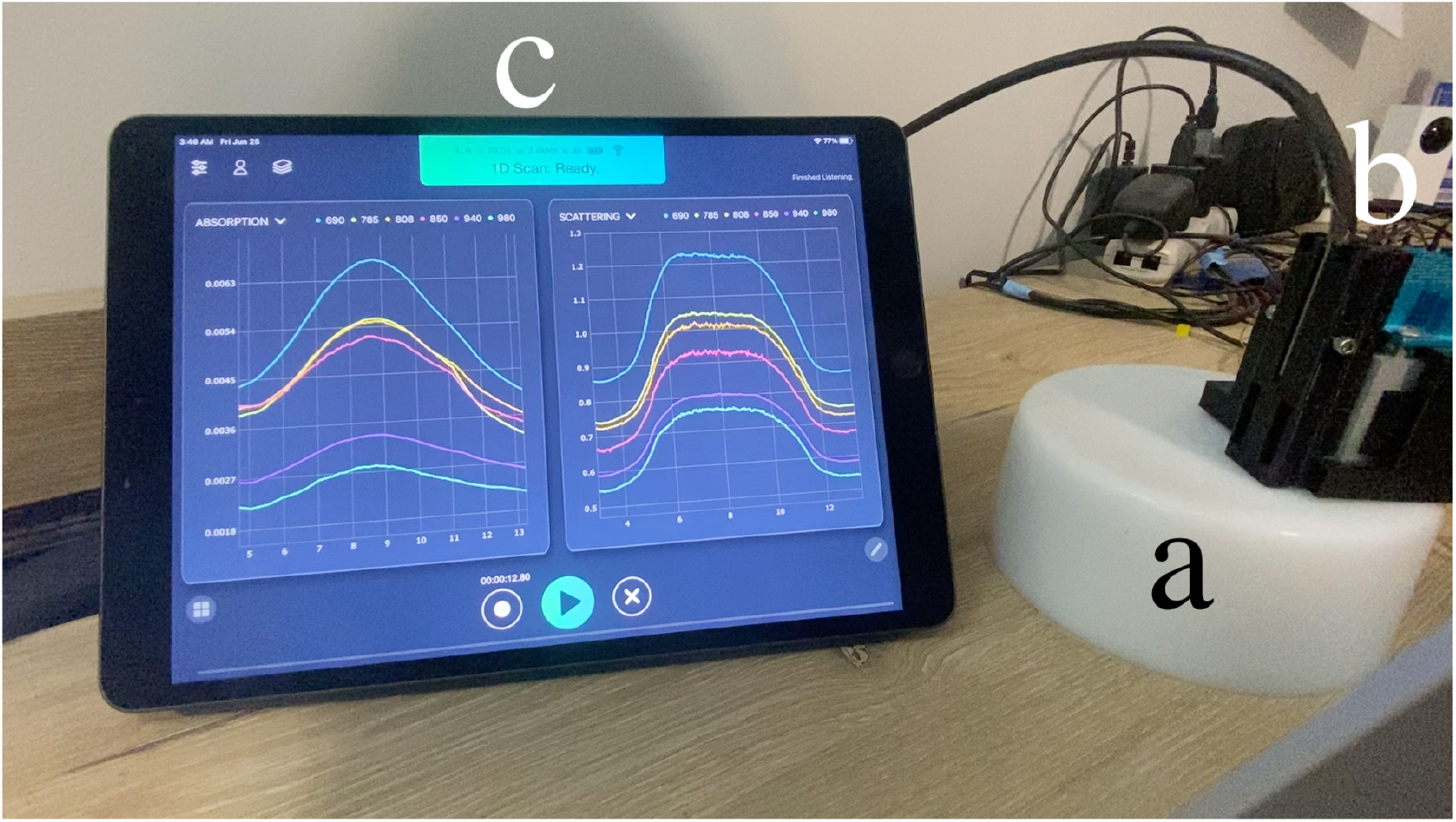
(a) A tissue simulating phantom (designed for *μ_a_*= 0.006 *mm*^−1^, *μ_s_*= 0.9 *mm*^−1^ at 660 nm) is impregnated with a 3.0 cm simulated tumor inclusion (designed for *μ_a_*= 0.01 *mm*^−1^, *μ_s_*= 1.3 *mm*^−1^ at 660 nm) 1 mm below the surface. (b) Six sources are fiber coupled to a handheld probe housing a 3 mm APD detector. (c) Absorption and reduced scattering values from the system are wirelessly transferred and displayed in real time at high speed (2,328 Hz for all six wavelengths) to a mobile device. The simulated tumor optical properties are visualized on the display as the probe is moved along the surface over the top of the tumor (video available with online version).

**Fig. 7.**
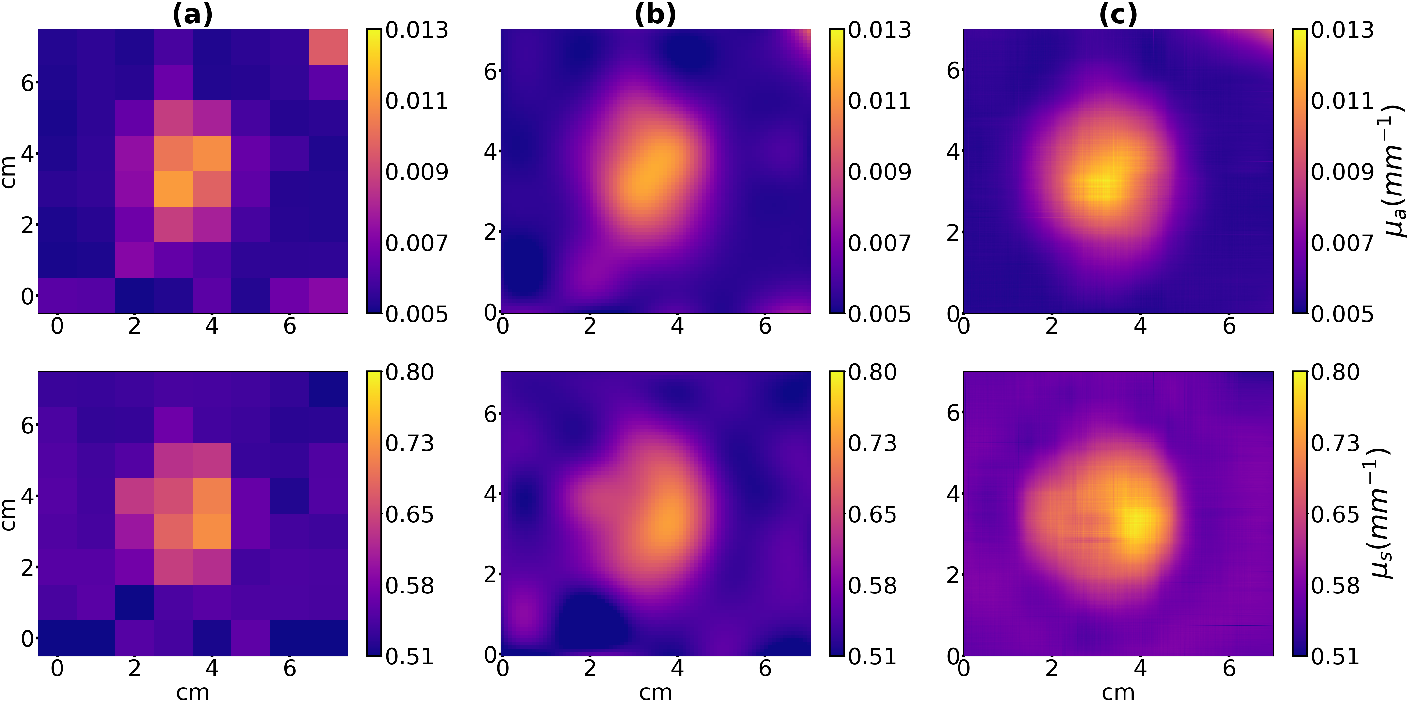
**(a)** A 7 × 7 cm image of a tissue simulating phantom with raw data at 1 × 1 cm spacing (a typical spatial measurement for FD-NIRS systems) comprising 49 points. **(b)** Bi-cubic interpolation of (a). **(c)** High-density 2D spatial maps acquired with the system at 352,000 points per image, 2,112,000 total samples (for all six wavelengths), bi-cubic interpolation, 3 cm diameter inclusion at 850 nm and 70 MHz modulation frequency. The high-density image more accurately recovers the 3 cm inclusion shape.

### N. In Vivo Real-time Display

To demonstrate the real-time processing and visualization abilities of the system, we performed an in vivo arterial occlusion measurement using a blood pressure cuff placed on the upper arm of a healthy volunteer. Amplitude and phase data taken with the system were recorded at 2.6 kHz using all 6 sources. Fig. 8 shows *HbO*_2_ and *HHb* concentrations recorded during the experiment. The data is displayed with a 1000-point (0.38s) moving average plotted for each chromophore. Cuff occlusion began at the 90 second mark, after which we observe a slight increase in *HbO*_2_, likely due to slow manual cuff inflation. The data then shows a steady falling concentration of *HbO*_2_, while *HHb* concentration rises as expected due to the arterial occlusion. After 5 minutes of occlusion, the cuff was released and a sharp hyperemic response and rebound was observed.

**Fig. 8.**
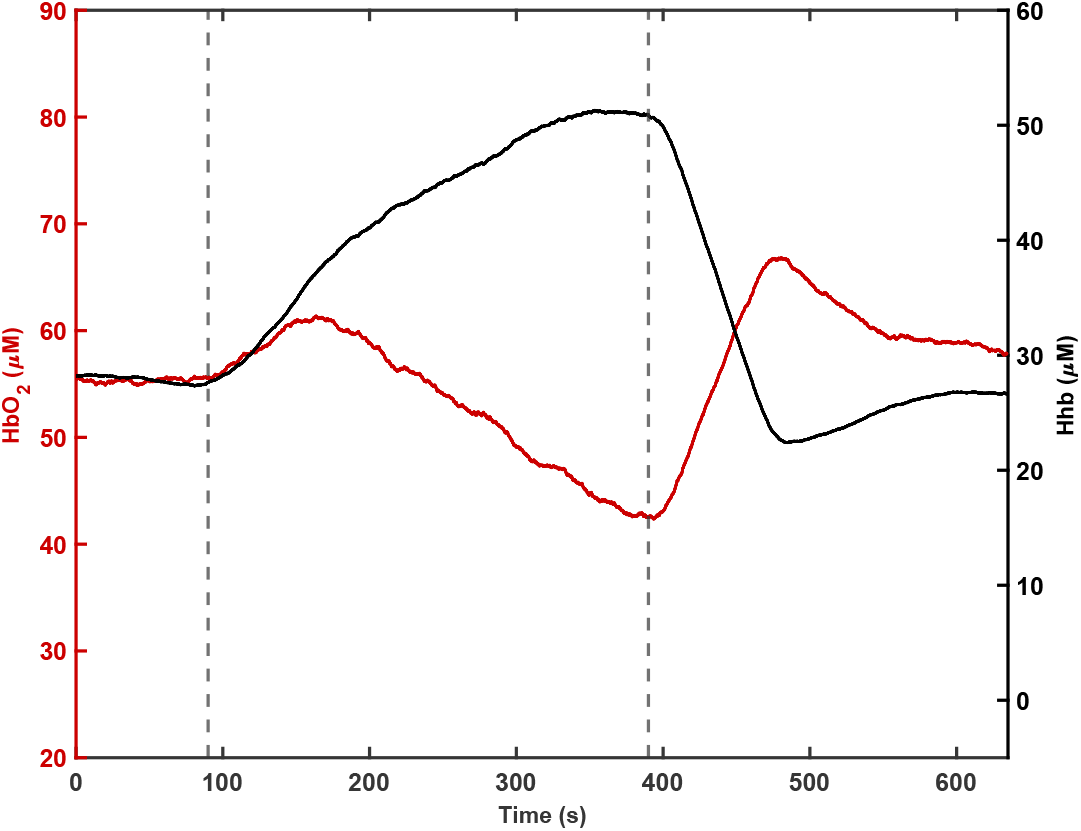
An arterial occlusion experiment showing tissue *HHb* (right) and *HbO*_2_ (left) curves from the forearm of a healthy human subject. Lines indicate the start and end of the occlusion on the upper arm. Measurements were acquired at 2.6 kHz with a 1000-point moving average plotted for each chromophore.

In a demonstration of the the system’s extremely high-speed sampling, measurements of chromophore data from two wavelengths (689 nm and 850 nm) on a healthy volunteer’s wrist are shown in Fig. 9. A pulsatile signal is clearly shown in the *HbO*_2_ curve and was measured with the systems to be 67 beats-per-minute (BPM), matching manual palpation using a stopwatch. The majority of the high temporal resolution pulse waveforms show the dicrotic notch, and we also observe the respiratory envelope superimposed on the *HbO*_2_ curve, estimated to be just under 10 seconds per breath. The inset in Fig. 9 displays *HbO*_2_ changes displayed in a zoomed curve of a single pulse peak where high-density points are indicated by red markers.

**Fig. 9.**
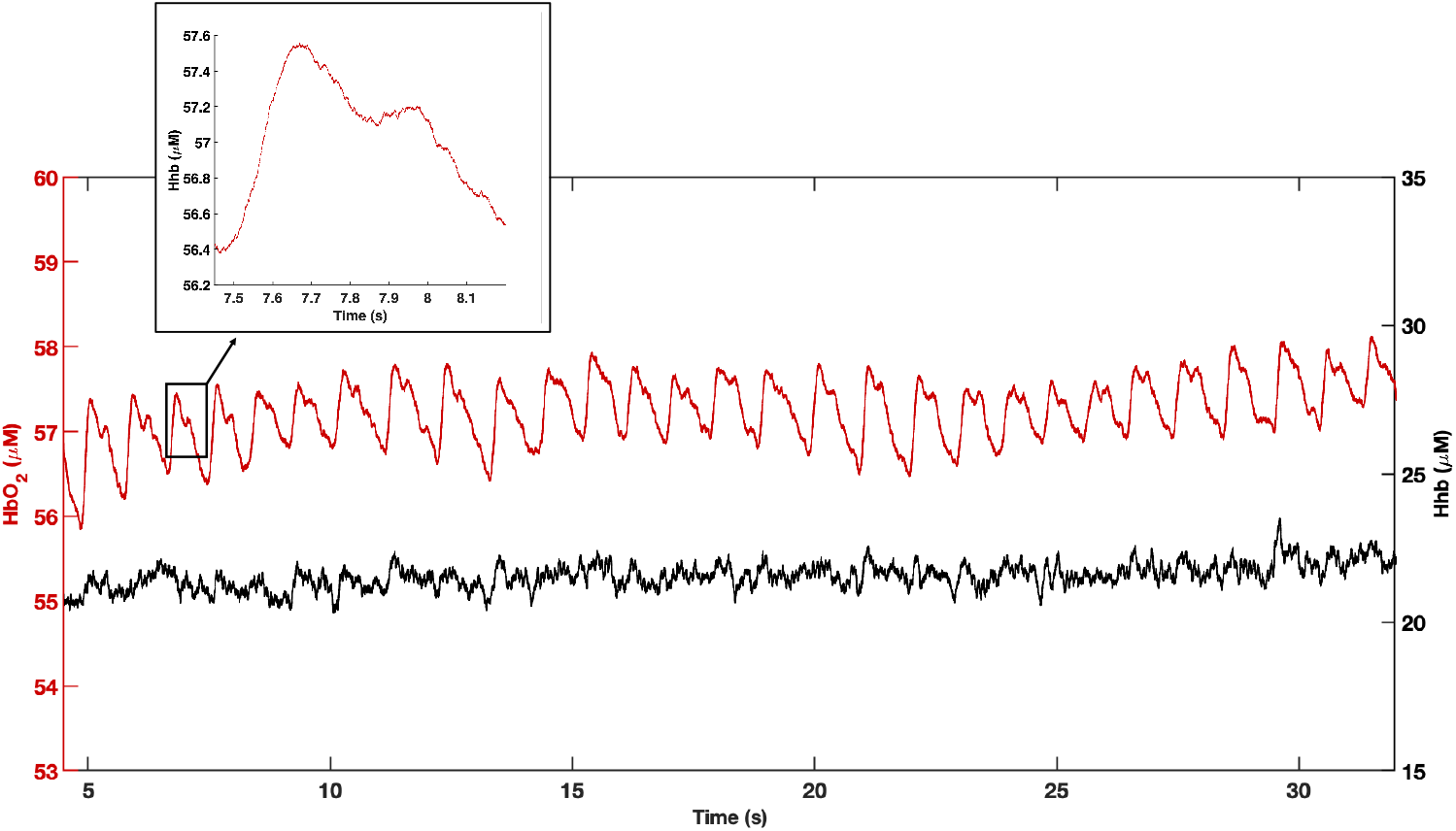
A pulsatile signal (~67 BPM) from a human wrist sampled at an extremely high-rate (5.4 kHz) is shown. The *HbO*_2_ curve displays a respiratory envelope function due to normal subject breathing (~10 second cycles). Inset: *HbO*_2_ changes are displayed in a zoomed curve of a single pulse peak where high-density points are indicated by red markers.

## Discussion and Conclusion

Over 25 years of clinical research has shown that FD-NIRS based optical imaging can be a meaningful tool for understanding, diagnosing, and treating various human diseases. For example, the ACRIN 6691 multi-center study demonstrated that FD-NIRS based methods predict neoadjuvant chemotherapy response in breast cancer with performance similar or better than MRI (34). For functional near-infrared spectroscopy (fNIRS), FD-NIRS can provide deeper sensitivity as compared to CW-fNIRS and therefore higher sensitivity to functional activation (31). FD-NIRS has also shown promising clinical utility in breast cancer diagnosis, critical care, exercise physiology, among other applications. However, there is currently only one company producing FD-NIRS systems (ISS, Inc.), and none are FDA-cleared for clinical use. Given our experience, we suggest that there are three major limitations that have contributed to the slow clinical adoption of FD-NIRS. First, all diffuse imaging methods typically exhibit low spatial resolution, thus providing data unlike standard medical imaging modalities. Second, the FD-NIRS research tools developed so far have been complex, slow, and prohibitively difficult to use. Finally, previous designs have lacked easy scalability necessary for high density and higher resolution imaging without greatly increasing the instrumentation footprint. The FD-NIRS platform we developed in this work was designed to overcome these outstanding issues through improved scalability, ease of use, and speed. Together, we believe that these improvements can lead to the higher spatial resolution, localization accuracy, sensitivity, and usability needed to accelerate clinical adoption.

Thus we demonstrated an all digital, scalable FD-NIRS platform capable of ultra-high speed chromophore processing and display utilizing an embedded system and external display. Digital ICs, RF and DC switches, filters and dynamic signal chain components were all specifically chosen for scalability, high-speed performance, and low noise. The DDS and RF circuitry allows for single or multi-frequency direct modulation of laser sources similar to VNA-based counterparts (1). This design can enable, for example, a fully embedded FD-NIRS system in a single handheld device, which could increase FD-NIRS portability and accessibility. The system also showed comparable noise levels to VNA-based systems (1, 35). We characterized the accuracy of the system in comparison to a reference system and found it to be within 10% across all wavelengths, suggesting acceptable equivalence.

We showed that FD-NIRS amplitude/phase, optical properties, and chromophores can be acquired, processed entirely on an embedded platform, and displayed at speeds up to 36,608 Hz, 17,891 Hz, and 10,211 Hz respectively. To the best of our knowledge, this is the fastest FD-NIRS system demonstrated to date, especially in its capability to calculate and display optical properties. For comparison, Zhao et al. showed DNN based chromophore recovery and display at rates up to 27 Hz (36), Applegate et al. streamed FD-NIRS data to a display at 1.5 Hz (33), and the ISS OxiplexTS200 and ISS Imagent report 100 Hz rates. In comparison, the platform presented here can display optical property and chromophore data between 100x to 3,600x faster than these systems, as illustrated in Table 2. Our platform includes two key innovations to achieve this performance. The FPGA-based Goertzel implementation enabled amplitude and phase information to be extracted from the raw ADC time domain signal at just 1/2 of the actual sampling speed (first demonstrated in (37, 38)). This strategy allows for a dramatic reduction in data storage (37,000x for 65,536 samples) and is a fundamental requirement to streaming data in real-time due to the large size of raw sampled ADC data. Second, we implemented an ultra-high-speed knn lookup table method into hardware, further reducing downstream computation and maximizing inversion/s speed, while also allowing chromophore computation to be done within the embedded SoC. Our knn lookup table method resulted in a 12 - 1,900x (in SoC) and 170 - 11,000x (modern CPU) improvement over previously reported literature (39). The lookup table approach was chosen as it can be significantly faster (1,000x) than iterative linear optimization solutions typically used to find inverse model solutions without loss in accuracy (17). Deep neural network (DNN) approaches can also provide fast inversion solutions (200 Hz), though at the expense of larger average errors than typical iterative solutions (36). DNN solutions are also limited by their sensitivity to input parameters; a new DNN model may be required for each SDS, modulation frequency and may be more susceptible to noise (36). A valid question to ask regarding high speed NIRS systems such as these is: why is it necessary to go so fast? We believe that these extremely high speeds provide significant advantages in scalability, accuracy, and imaging resolution. First, there is a desire among fNIRS researchers and users to cover the human head with greater area and density of optodes. It is estimated that 370 optodes are needed to cover the adult scalp for high-density DOT imaging (40). Our platform can easily scale to this number of optodes and thus provide quantitative, physiological information over the entire scalp in real-time. This could be of major clinical significance for patients that lack baseline measurements, such as those with brain trauma. The increased speed and sensitivity could also be used to further investigate the fast (<100ms) optical scattering-driven evoked-response optical signals that may be indicative of neural activity (41, 42), but difficult to measure.

Second, high-speed FD-NIRS acquisition and processing gives the user significant control over modulation frequency choices and data averaging. These parameters can be optimized specifically for the desired application all while providing real-time display, similar to the various tissue modes of an ultrasound device. For example, some applications and tissue types can benefit from a range of modulation frequencies, while others require only one (30, 39). We also showed that the large amount of streaming data can be averaged to provide an order of magnitude improvement in phase precision at the expense of speed if desired.

Finally, this real-time high-speed FD-NIRS platform can be used to provide higher spatial density imaging of the tissue and lead to better assessment of tissue heterogeneity and overall resolution. To illustrate, we generated ultra-high-spatial-density 1D and 2D scans of phantoms with tumor-simulating inclusions at sizes from 1 - 3 cm. The results reveal a smooth, information-rich assessment of the phantom’s optical properties, as well as the ability to accurately differentiate the inclusion sizes. A 7 cm line scan in Fig. 5 took 12 seconds, demonstrating significantly faster imaging speed than other quantitative NIRS approaches (12, 33). The 2D quantitative optical property topography maps (Fig.7) contain 352,000 points per image, representing 2,112,000 samples (for 6*λ*). This represents highest density 2D topographical FD-NIRS image created to date, with 43,100x times more spatial density than the grids used in ACRIN 6691 (12), and chromophore display speeds 6,800x faster than previous works (33). Accurately assessing spatial heterogeneity can be significant in clinical applications; for example, it may be a significant marker in determining neoadjuvant chemotherapy response (43). Importantly, the images can be constructed in real-time, allowing a user to re-take data if the acquisition was poor or to dynamically adjust the imaging region of interest. In conclusion, we have demonstrated a new FD-NIRS platform with unprecedented scalability and speed that can lead to new compact and potentially wearable quantitative diffuse optical spectroscopy devices. These improvements are not only applicable to existing applications, such as fNIRS and breast cancer imaging, but can also open up new applications as the technology is further disseminated. Our future work entails miniaturization of the platform into handheld and wearable form factors, as well as application of the platform to acquire high resolution 2D and 3D diffuse optical imaging of tissue.

## Disclosures

RAS, VJK, and TDO disclose pending and granted patents related to FD-NIRS technology. RAS, AC, LS, and TDO disclose ownership of NearWave Corp., which is producing FD-NIRS technologies.

## Acknowledgements

This work was funded by the University of Notre Dame, Harper Cancer Research Institute’s Research Like a Champion program, and the IDEA Center at Notre Dame.

## Bibliography

1. Bruce J. Tromberg, Natasha Shah, Ryan Lanning, Albert Cerussi, Jennifer Espinoza, Tuan Pham, Lars Svaasand, and John Butler. Non-invasive in vivo characterization of breast tumors using photon migration spectroscopy. Neoplasia, 2(1-2):26–40, 2000. ISSN 15228002. doi: 10.1038/sj.neo.7900082.

2. B. Chance, M. Cope, E. Gratton, N. Ramanujam, and B. Tromberg. Phase measurement of light absorption and scatter in human tissue. Review of Scientific Instruments, 69(10):3457–3481, 1998. ISSN 00346748. doi: 10.1063/1.1149123.

3. Frédéric Bevilacqua, Dominique Piguet, Pierre Marquet, Jeffrey D. Gross, Bruce J. Tromberg, and Christian Depeursinge. In vivo local determination of tissue optical properties: applications to human brain. Applied Optics, 38(22):4939, 1999. ISSN 0003-6935. doi: 10.1364/ao.38.004939.

4. L. Meng, A. W. Gelb, B. S. Alexander, A. E. Cerussi, B. J. Tromberg, Z. Yu, and W. W. Mantulin. Impact of phenylephrine administration on cerebral tissue oxygen saturation and blood volume is modulated by carbon dioxide in anaesthetized patients. British Journal of Anaesthesia, 108(5):815–822, 2012. ISSN 14716771. doi: 10.1093/bja/aes023.

5. Albert Cerussi, Richard Van Woerkom, Feizal Waffarn, and Bruce Tromberg. Noninvasive monitoring of red blood cell transfusion in very low birthweight infants using diffuse optical spectroscopy. Journal of Biomedical Optics, 10(5):051401, 2005. ISSN 10833668. doi: 10.1117/1.2080102.

6. Thomas D O’Sullivan, Anaïs Leproux, Jeon-Hor Chen, Shadfar Bahri, Alex Matlock, Darren Roblyer, Christine E McLaren, Wen-Pin Chen, Albert E Cerussi, Min-Ying Su, and Bruce J Tromberg. Optical imaging correlates with magnetic resonance imaging breast density and revealscomposition changes during neoadjuvant chemotherapy. Breast Cancer Research, 15(1):R14, 2013. ISSN 1465-542X. doi: 10.1186/bcr3389.

7. Goutham Ganesan, Joshua A Cotter, Warren Reuland, Albert E Cerussi, Bruce J Tromberg, and Pietro Galassetti. Effect of blood flow restriction on tissue oxygenation during knee extension. Medicine and science in sports and exercise, 47(1):185–193, jan 2015. ISSN 1530-0315. doi: 10.1249/MSS.0000000000000393.

8. Tuan H Pham, Renee Hornung, Hongphuc P Ha, Tanya Burney, Dan Serna, Ledford Powell, Matthew Brenner, and Bruce J Tromberg. Non-invasive monitoring of hemodynamic stress using quantitative near-infrared frequency-domain photon migration spectroscopy. Journal of Biomedical Optics, 7(1):34–44, 2002. doi: 10.1117/1.1427046.

9. Matthaios Doulgerakis, Adam T Eggebrecht, and Hamid Dehghani. High-density functional diffuse optical tomography based on frequency-domain measurements improves image quality and spatial resolution. Neurophotonics, 6(3):1–14, 2019. doi: 10.1117/1.NPh.6.3.035007.

10. Thomas D. O’Sullivan, Albert E. Cerussi, David J. Cuccia, and Bruce J. Tromberg. Diffuse optical imaging using spatially and temporally modulated light. Journal of Biomedical Optics, 17(7): 0713111, 2012. ISSN 1083-3668. doi: 10.1117/1.JBO.17.7.071311.

11. Anaïs Leproux, Thomas D. O’Sullivan, Albert Cerussi, Amanda Durkin, Brian Hill, Nola Hylton, Arjun G. Yodh, Stefan A. Carp, David Boas, Shudong Jiang, Keith D. Paulsen, Brian Pogue, Darren Roblyer, Wei Yang, and Bruce J. Tromberg. Performance assessment of diffuse optical spectroscopic imaging instruments in a 2-year multicenter breast cancer trial. Journal of Biomedical Optics, 22(12):1, 2017. ISSN 1560-2281. doi: 10.1117/1.jbo.22.12.121604.

12. Bruce J. Tromberg, Zheng Zhang, Anaïs Leproux, Thomas D. O’Sullivan, Albert E. Cerussi, Philip M. Carpenter, Rita S. Mehta, Darren Roblyer, Wei Yang, Keith D. Paulsen, Brian W. Pogue, Shudong Jiang, Peter A. Kaufman, Arjun G. Yodh, So Hyun Chung, Mitchell Schnall, Bradley S. Snyder, Nola Hylton, David A. Boas, Stefan A. Carp, Steven J. Isakoff, and David Mankoff. Predicting responses to neoadjuvant chemotherapy in breast cancer: ACRIN 6691 trial of diffuse optical spectroscopic imaging. Cancer Research, 76(20):5933–5944, 2016. ISSN 15387445. doi: 10.1158/0008-5472.CAN-16-0346.

13. G. Ganesan, R. V. Warren, A. Leproux, M. Compton, K. Cutler, S. Wittkopp, G. Tran, T. O’Sullivan, S. Malik, P. R. Galassetti, and B. J. Tromberg. Diffuse optical spectroscopic imaging of subcutaneous adipose tissue metabolic changes during weight loss. International Journal of Obesity, 40(8):1292–1300, 2016. ISSN 14765497. doi: 10.1038/ijo.2016.43.

14. Alyssa Torjesen, Raeef Istfan, and Darren Roblyer. Ultrafast wavelength multiplexed broad bandwidth digital diffuse optical spectroscopy for in vivo extraction of tissue optical properties. Journal of Biomedical Optics, 22(3):036009, 2017. ISSN 1083-3668. doi: 10.1117/1.JBO.22.3.036009.

15. Bernhard B. Zimmermann, Qianqian Fang, David A. Boas, and Stefan A. Carp. Frequency domain near-infrared multiwavelength imager design using high-speed, direct analog-to-digital conversion. Journal of Biomedical Optics, 21(1):016010, 2016. ISSN 1083-3668. doi: 10.1117/1.JBO.21.1.016010.

16. Darren Roblyer, Thomas D. O’Sullivan, Robert V. Warren, and Bruce J. Tromberg. Feasibility of direct digital sampling for diffuse optical frequency domain spectroscopy in tissue. Measurement Science and Technology, 24(4), 2013. ISSN 13616501. doi: 10.1088/0957-0233/24/4/045501.

17. Tuan H. Pham, Olivier Coquoz, Joshua B. Fishkin, Eric Anderson, and Bruce J. Tromberg. Broad bandwidth frequency domain instrument for quantitative tissue optical spectroscopy. Review of Scientific Instruments, 71(6):2500–2513, 2000. ISSN 00346748. doi: 10.1063/1.1150665.

18. N S Altman. An Introduction to Kernel and Nearest-Neighbor Nonparametric Regression. The American Statistician, 46(3):175–185, 1992. doi: 10.1080/00031305.1992.10475879.

19. Petr Sysel and Pavel Rajmic. Goertzel algorithm generalized to non-integer multiples of fundamental frequency. EURASIP Journal on Advances in Signal Processing, 2012(1):56, mar 2012. ISSN 1687-6180. doi: 10.1186/1687-6180-2012-56.

20. Alyssa Torjesen, Raeef Istfan, Hannah Peterson, and Darren Roblyer. An Ultra-Fast Digital Diffuse Optical Spectroscopic Imaging (dDOSI) System for Monitoring Chemotherapy Response. Biomedical Optics 2016, 2016:OTh3C.1, 2016. doi: 10.1364/OTS.2016.OTh3C.1.

21. Arjun Yodh and Britton Chance. Spectroscopy and Imaging with Diffusing Light. Physics Today, 48(3):34–40, 1995. ISSN 19450699. doi: 10.1063/1.881445.

22. Irving Bigio. Quantitative biomedical optics : theory, methods, and applications. Cambridge University Press, Cambridge, 2016. ISBN 9780521876568.

23. W. G. Zijlstra and A. Buursma. Spectrophotometry of hemoglobin: Absorption spectra of bovine oxyhemoglobin, deoxyhemoglobin, carboxyhemoglobin, and methemoglobin. Comparative Biochemistry and Physiology - B Biochemistry and Molecular Biology, 118(4):743–749, 1997. ISSN 03050491. doi: 10.1016/S0305-0491(97)00230-7.

24. A. E. Cerussi, D. Jakubowski, N. Shah, F. Bevilacqua, R. Lanning, A. J. Berger, D. Hsiang, J. Butler, R. F. Holcombe, and B. J. Tromberg. Spectroscopy enhances the information content of optical mammography. Journal of Biomedical Optics, 7(1):60, 2002. ISSN 10833668. doi: 10.1117/1.1427050.

25. R. L. P. van Veen, H. J. C. M. Sterenborg, A. Pifferi, A. Torricelli, E. Chikoidze, and R. Cubeddu. Determination of visible near-IR absorption coefficients of mammalian fat using time- and spatially resolved diffuse reflectance and transmission spectroscopy. Journal of Biomedical Optics, 10(5):054004, 2005. ISSN 10833668. doi: 10.1117/1.2085149.

26. W.R. Bennett. pectra of Quantized Signals. Bell System Technical Journal, 27:446–471, jul 1948.

27. Albert E. Cerussi, Robert Warren, Brian Hill, Darren Roblyer, Anais Leproux, Amanda F. Durkin, Thomas D. O’Sullivan, Sam Keene, Hosain Haghany, Timothy Quang, William M. Mantulin, and Bruce J. Tromberg. Tissue phantoms in multicenter clinical trials for diffuse optical technologies. Biomedical Optics Express, 3(5):966, 2012. ISSN 2156-7085. doi: 10.1364/boe.3.000966.

28. Sergio Fantini and Angelo Sassaroli. Frequency-Domain Techniques for Cerebral and Functional Near-Infrared Spectroscopy. Frontiers in Neuroscience, 14(April):1–18, 2020. ISSN 1662453X. doi: 10.3389/fnins.2020.00300.

29. Grant Fowles. Introduction to modern optics. Dover Publications, New York, 1989. ISBN 0486659577.

30. Guy A. Perkins, Adam T. Eggebrecht, and Hamid Dehghani. Quantitative evaluation of frequency domain measurements in high density diffuse optical tomography. Journal of Biomedical Optics, 26, 2021. ISSN 15602281. doi: 10.1117/1.jbo.26.5.056001.

31. Matthaios Doulgerakis, Adam T. Eggebrecht, and Hamid Dehghani. High-density functional diffuse optical tomography based on frequency-domain measurements improves image quality and spatial resolution. Neurophotonics, 6(03):1, 2019. ISSN 2329-4248. doi: 10.1117/1.nph.6.3.035007.

32. Ola Abdalsalam, Yide Zhang, Scott Howard, and Thomas O’Sullivan. Self-calibrated frequency domain diffuse optical spectroscopy with a phased source array. In Sergio Fantini, Bruce J Tromberg, Eva Marie Sevick-Muraca, and Paola Taroni, editors, Optical Tomography and Spectroscopy of Tissue XIII, volume 10874, pages 1–6. International Society for Optics and Photonics, SPIE, 2019. doi: 10.1117/12.2510422.

33. Matthew Applegate, Robert Amelard, Carlos Gomez, and Darren Roblyer. Real-time handheld probe tracking and image formation using digital frequency-domain diffuse optical spectroscopy. IEEE Transactions on Biomedical Engineering, 2021. ISSN 15582531. doi: 10.1109/TBME.2021.3072036.

34. Jeffrey M. Cochran, David R. Busch, Anaïs Leproux, Zheng Zhang, Thomas D. O’Sullivan, Albert E. Cerussi, Philip M. Carpenter, Rita S. Mehta, Darren Roblyer, Wei Yang, Keith D. Paulsen, Brian Pogue, Shudong Jiang, Peter A. Kaufman, So Hyun Chung, Mitchell Schnall, Bradley S. Snyder, Nola Hylton, Stefan A. Carp, Steven J. Isakoff, David Mankoff, Bruce J. Tromberg, and Arjun G. Yodh. Tissue oxygen saturation predicts response to breast cancer neoadjuvant chemotherapy within 10 days of treatment. Journal of Biomedical Optics, 24(02):1, 2018. ISSN 1560-2281. doi: 10.1117/1.jbo.24.2.021202.

35. SJ Madsen, ER Anderson, RC Haskell, and BJ Tromberg. frequency-domain photon migration, optical properties, diode laser, network analyzer. In Proc. SPIE, volume 2389, pages 257–263, 1995.

36. Yanyu Zhao, Mattew B Applegate, Raeef Istfan, Ashvin Pande, and Darren Roblyer. Quantitative real-time pulse oximetry with ultrafast frequency-domain diffuse optics and deep neural network processing. Biomedical Optics Express, 9(12):5997–6008, 2018. doi: 10.1364/BOE.9.005997.

37. Vincent J Kitsmiller, Roy A Stillwell, and Thomas D O’Sullivan. Toward handheld real time frequency domain diffuse optical spectroscopy. In Sergio Fantini, Bruce J Tromberg, Eva Marie Sevick-Muraca, and Paola Taroni, editors, Optical Tomography and Spectroscopy of Tissue XIII, volume 10874, pages 13–19. International Society for Optics and Photonics, SPIE, 2019.

38. Roy A. Stillwell, Vincent J. Kitsmiller, and Thomas D. O’Sullivan. Towards a high-speed handheld frequency-domain diffuse optical spectroscopy deep tissue imaging system. In Biophotonics Congress: Biomedical Optics 2020 (Translational, Microscopy, OCT, OTS, BRAIN), page TTu1B.7. Optical Society of America, 2020. doi: 10.1364/TRANSLATIONAL.2020.TTu1B.7.

39. Matthew B Applegate, Carlos A Gómez, and Darren M. Roblyer. Modulation frequency selection and efficient look-up table inversion for frequency domain diffuse optical spectroscopy. Journal of Biomedical Optics, 26(03):1–15, 2021. ISSN 1560-2281. doi: 10.1117/1.jbo.26.3.036007.

40. Hubin Zhao and Robert J Cooper. Review of recent progress toward a fiberless, whole-scalp diffuse optical tomography system. Neurophotonics, 5(01):1, 2017. ISSN 2329-423X. doi: 10.1117/1.nph.5.1.011012.

41. Gabriele Gratton and Monica Fabiani. Fast optical imaging of human brain function. Frontiers in Human Neuroscience, 4:52, 2010. ISSN 1662-5161. doi: 10.3389/fnhum.2010.00052.

42. Gabriele Gratton, Paul M Corballis, Eunhee Cho, Monica Fabiani, and Donald C Hood. Shades of gray matter: Noninvasive optical images of human brain reponses during visual stimulation. Psychophysiology, 32(5):505–509, 1995. doi: https://doi.org/10.1111/j.1469-8986.1995.tb02102.x.

43. Ylenia Santoro, Bruce J. Tromberg, Enrico Gratton, Anais Leproux, and Albert E. Cerussi. Breast cancer spatial heterogeneity in near-infrared spectra and the prediction of neoadjuvant chemotherapy response. Journal of Biomedical Optics, 16(9):097007, 2011. ISSN 10833668. doi: 10.1117/1.3638135.

