## Supplementary Data for "A scalable, multi-wavelength, broad bandwidth frequency-domain near-infrared spectroscopy platform for real-time quantitative tissue optical imaging"

Fig.S1 shows high-density 2D spatial optical property maps with 2,112,000 measurements (352,000 points per image) made from a tissue simulating phantom with a buried inclusion showing $\mu_a$, $\mu_s$ data for six different sources as explained in the main text. A probe is scanned across the phantom surface in the horizontal direction, then in the vertical direction in a 7 cm x 7 cm rectangle. The image acquisition time is around 3 minutes total.

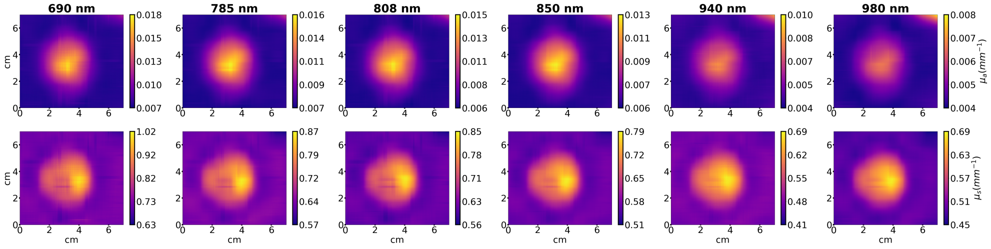


**Fig. S1.** High-density 2D spatial optical property maps made from a tissue simulating phantom with a buried inclusion showing $\mu_a$, $\mu_s$ data for six different sources.
